# SRAscout: Semantic discovery of public sequencing data through harmonized SRA and BioProject metadata

**DOI:** 10.64898/2026.11.15.766670

**Authors:** Dimitris Grigoriadis, Nikos Perdikopanis, Candace S.Y. Chan, Georgios A. Pavlopoulos, Ilias Georgakopoulos-Soares

## Abstract

The rapid growth of public sequencing data has created great opportunities for data-driven discovery in the life sciences, but it has also made it harder to find relevant datasets. The NCBI Sequence Read Archive (SRA) hosts more than 45 million high-throughput sequencing runs, and the BioProject database, shared by NCBI, the European Nucleotide Archive (ENA) and the DNA Data Bank of Japan, describes the projects behind them. However, their metadata are often incomplete, inconsistently annotated and difficult to search using keywords alone. We present SRAscout, a web-based platform that enables semantic dataset retrieval by integrating SRA run metadata with the corresponding BioProject records. SRAscout collects and indexes metadata for about 714,000 BioProjects, and harmonizes sample attributes such as cell line, tissue, age, sex and treatment into a compact and consistent schema. For semantic embedding, SRAscout employs a biomedical sentence embedding model based on BioBERT, which generates dense vector representations from the title, name, organism and description of each project. Users can search in three modes, with the DOI or PubMed ID of an article, with free text, or with BioProject and SRA study accessions, and SRAscout finds the most related projects in each case. Free-text queries are processed through a hybrid of semantic and full-text search, which finds conceptually relevant studies even when different terms are used to describe the same biology, while exact terms such as gene names are still matched. Structured filters allow refinement by organism, assay type (for example RNA-seq and miRNA-seq), harmonized sample attributes such as tissue, cell type and disease, tag words, and many additional attributes. For each search, users can download a harmonized metadata table and ready-to-use scripts that fetch the sequencing files from ENA. SRAscout also offers a public API for programmatic access. By combining harmonized metadata with context-aware search in a single interface, SRAscout facilitates dataset discovery, meta-analysis and data reuse. The application is freely available at www.srascout.com.

## 1. Introduction

Public sequencing archives have changed the way biological research is done. The NCBI Sequence Read Archive (SRA) [1] and the European Nucleotide Archive (ENA) [2] are two of the three partners of the International Nucleotide Sequence Database Collaboration, and together they store raw data from a very large number of experiments. The SRA metadata snapshot of November 2026 alone contains 45.2 million public runs from about 714,000 BioProjects. This growth enables researchers to leverage publicly available datasets, compare their own findings with published studies, and generate novel hypotheses without the need to generate additional sequencing data. For example, public CAGE data have been used to train deep learning models that identify transcription start sites [3,4], and public transcriptomic data from many studies have been combined into resources of stress-induced transcriptional activity in human cell lines [5]. However, depositing data in an archive does not make it easy to find. Despite the FAIR principles (Findable, Accessible, Interoperable, Reusable) [6], discovering the datasets that best match a specific biological question remains challenging.

One of the main obstacles is the quality of the metadata. Sample descriptions are written by the submitters as free text, and field names and values are rarely standardized. A large study of the BioSample and BioSamples repositories showed that most metadata fields are not controlled and that the same property of a sample is often described in many different ways [7]. The same problem affects SRA, where the metadata contain many synonyms and spelling variants [8]. As a result, keyword search only finds studies that use the same words as the query, and it misses studies that describe the same biology in different terms.

Several tools were created to make SRA metadata easier to use. SRAdb [9] and pysradb [10] allow users to query SRA metadata from R and Python, while iSeq [11] and the nf-core/fetchngs pipeline [12] retrieve metadata and sequencing files for accessions that the user already knows. SRA-Explorer [13] offers a simple web page to find runs and get download links from SRA and ENA and DDBJ Search [14] provides keyword search over the BioProject, BioSample and SRA records of all three INSDC partners. MetaSRA [8] normalized the metadata of human samples using biomedical ontologies, and MetaHQ [15] has recently extended the harmonization of metadata to public omics samples and studies. Large language models have also been applied to this problem. SRAgent [16] uses LLM agents to find single-cell datasets through the NCBI Entrez utilities and to extract structured metadata for each experiment, while SRAKE [17] indexes SRA metadata locally and adds vector search over study descriptions in a command-line tool. Other resources focus on the data itself. recount3 [18] and ARCHS4 [19] provide uniformly processed RNA-seq data for human and mouse, and the Omics Discovery Index [20] links datasets from several omics repositories. Logan-Search [21] and MetaGraph [22] search the sequences of SRA runs directly, which answers a different question from metadata search These resources are very useful, but they require the user to know the accessions in advance, either focus on selected organisms or data types, or they still depend on keyword search over the original text, or run as local software rather than as a public web service.

Language models offer a different path. Domain-specific models such as BioBERT [23], and sentence-level models built on the Sentence-BERT approach [24], convert text into numeric vectors so that texts with similar meaning end up close to each other, even when they use different words. This idea was recently tested on GEO by Brown and colleagues, who showed that embeddings of dataset descriptions often find relevant series more effectively than the GEO search engine, and who released a web application called GEOfinder [25]. GEOfinder starts from datasets that the user already knows and looks for similar ones. In our previous work, we developed Public Omics Explorer (POE) [26], which accepts natural language queries and links GEO datasets and ENA records through their PubMed publications. Because POE relies on the text of linked articles, it covers only datasets that have a publication in PubMed, and it does not reach sequencing projects that were deposited directly in SRA or ENA.

This leaves an important gap. In the current SRA snapshot, 75% of the BioProjects behind public runs have no linked publication and 73% have no GEO series, so neither publication-based nor GEO-based search can reach them. To our knowledge, no public web server lets researchers search the BioProjects behind all SRA runs with natural language, across all organisms and assay types, and then narrow the results with structured filters on harmonized sample attributes such as tissue, cell type, cell line, age, sex and treatment.

In this study, we present SRAscout, a free web server for the semantic discovery of public sequencing studies. SRAscout combines run-level metadata from NCBI SRA with the corresponding records of the BioProject database [27], which is shared by NCBI, ENA and the DNA Data Bank of Japan, and links each study to its sequencing files in ENA. Sample attributes such as cell line, tissue, age, sex and treatment are cleaned and mapped to a small and consistent set of fields. The title, name, organism and description of about 714,000 BioProjects are converted into embeddings with a biomedical sentence embedding model based on BioBERT [23]. Users can search in three modes. They can enter the DOI or PubMed ID of an article, whose title and abstract are retrieved automatically, write free text from a few words to a full paragraph, or enter BioProject and SRA study accessions, and in each case SRAscout finds the most related projects. For free text, a semantic search and a full-text search run together and their results are merged, so studies described in different words are found while exact terms such as gene names are still matched. Results can then be narrowed with filters on organism, assay type (for example RNA-seq and miRNA-seq), harmonized sample attributes, tag words and many other attributes, and users can download a harmonized metadata table and scripts that fetch the sequencing files. SRAscout needs no login and is freely available at www.srascout.com.

To evaluate SRAscout, we built a benchmark of 120 queries, and judged the relevance of the pooled results of four search methods, covering human and model organisms, microbiomes and environmental samples, and assay types such as RNA-seq, ChIP-seq and amplicon sequencing. On this benchmark, the hybrid search retrieved 60.3% of the relevant projects among the top 20 results, compared with 56.1% for semantic search alone, 30.6% for full-text search alone and 23.6% for the keyword search of the NCBI BioProject database. Semantic search was most helpful for queries that describe a topic in words that differ from those used by the submitters, while full-text search contributed exact identifiers such as gene names and strains.

## 2. Materials and Methods

### 2.1 Data sources and collection

SRAscout is built from three public sources. Run-level metadata come from the NCBI Sequence Read Archive (SRA) [1], project-level records come from the BioProject database [27], and sequencing file information comes from the European Nucleotide Archive (ENA) [2]. The BioProject database is shared by NCBI, ENA and DDBJ.

SRA run metadata are downloaded from the public NCBI SRA metadata bucket on Amazon S3, where they are stored in Apache Parquet format. The files are loaded into a local DuckDB database and exported as one table with a row for every run. As of November 2026, the snapshot contains 45.2 million runs. Each run record has the run, experiment, study, sample and BioSample accessions, the BioProject accession, organism, assay type, library name, library layout, library selection, library source, sequencing platform, instrument, sequencing center, sample country and release date. It also has a JSON field with the free-form attributes that submitters attached to the sample, experiment and run. These metadata are cleaned as described in Section 2.2. Runs without a BioProject accession (about 1.3%) are not used for project-level records.

Project records are taken from the BioProject XML file that NCBI publishes on its FTP server and updates daily. The file is approximately 4 GB, and it is read as a stream and only the projects that are referenced by SRA runs are kept. For each project SRAscout stores the name, title and description, the organism and taxonomy identifier, the project type and data types, the target material, the submission and release dates, the submitting organization and its country, grants, links to GEO accessions, parent projects, and linked publications with their PubMed identifiers, journals and dates. The records are organized as one row for each pair of BioProject and SRA study, covering 714,315 distinct BioProjects

Information about sequencing files is not copied into the SRAscout database. When a user opens a study, the file names, sizes, MD5 checksums and FASTQ download links are requested from the ENA Portal API by the user’s browser, so that ENA’s request limits apply to each user and not to our server.

The collection is refreshed every week by an automated process that downloads new records and updates the tables with new rows. The current release was built from the SRA metadata and BioProject file as of November 2026.

### 2.2 Metadata harmonization and normalization

Public metadata are written by thousands of different submitters, resulting in substantial variation in attribute names, terminology, and formatting. SRAscout cleans the metadata in two steps. First, the text of each BioProject record is cleaned and its fields are put in a consistent form. Second, the free form sample attributes of all SRA runs are mapped to a small set of fields and summarized for each BioProject.

#### 2.2.1 BioProject records

BioProject titles and descriptions often contain embedded HTML tags, encoded characters and irregular spacing. SRAscout removes the HTML tags, converts encoded characters back to plain text and collapses repeated whitespace, so that the text is clean for display, full text indexing and embedding. Controlled values are written in a readable form, dates are reduced to the day, and repeated values in multi valued fields are removed. Linked publications are parsed into structured citations with their identifier, title, journal, year and authors. GEO series accessions are collected from both the center identifiers and the external links of each project, so that projects that also exist in GEO can be recognized.

#### 2.2.2 Sample attributes

Every SRA run carries a JSON field with the attributes that the submitter attached to the sample, the experiment and the run. In the November 2026 snapshot, the 45.2 million runs used 99,295 distinct attribute keys. Many of these keys are versions of the same attribute with a suffix added by SRA, which shows where the attribute came from (for example *tissue_sam* and *tissue_run*) or separates repeated names. After these suffixes were removed, 95,582 distinct keys remained. Most of them were used by very few submitters. Only 145 keys appeared in at least 1% of runs, and 579 in at least 0.1% of runs.

To decide which information to keep, we counted how many runs use each key and grouped keys that describe the same property. A field was kept when its combined keys cover at least 5% of runs. The only exception is the molecular target, which covers about 1% of runs but is the only way to tell *ChIP-seq* experiments apart. This gave 14 fields (Table 1). Within each field, the source keys are listed in order of priority, with the most common and most specific key first, and for each run the first non-empty key is used. Keys that describe the host of a pathogen or microbiome sample (for example *host_tissue_sampled* or *host_sex*) were merged into the same fields, because for these projects the host is usually what a user is looking for. The host organism itself is kept as a separate field so that the two cases can still be told apart. Some frequent keys were left out on purpose. The isolate key appears in 34% of runs but is almost always a sample identifier and not a strain name, and keys about location and environment, such as collection date and coordinates, do not describe the biology of the sample. Phenotype was also left out, because its values were too varied to merge with disease state.

**Table 1.** Harmonized sample attribute fields, their source keys, and their coverage in the November 2026 snapshot (45.2 million runs, 714,315 BioProjects).

| Field | Source keys (in order of priority) | Runs with a value | BioProjects with a value |
|---|---|---|---|
| Organism | organism (run metadata) | 98.6% | 100.0% |
| Assay type | assay_type (run metadata) | 91.8% | 91.4% |
| Host | host, host_scientific_name, host_common_name | 36.0% | 15.7% |
| Tissue | tissue, organism_part, tissue_type, body_site, host_tissue_sampled, host_body_site | 28.6% | 35.4% |
| Sex | sex, host_sex, gender, host_gender | 25.1% | 13.0% |
| Genotype or strain | genotype, strain, cultivar, breed, ecotype, genetic_background, variety, host_genotype | 23.2% | 49.1% |
| Age | age, host_age | 18.6% | 14.1% |
| Source name | source_name | 17.1% | 23.1% |
| Disease state | disease, disease_state, disease_status, diagnosis, host_disease, host_health_state, health_state, host_disease_stat, health_status, clinical_status | 13.9% | 4.6% |
| Developmental stage | dev_stage, developmental_stage, development_stage, host_life_stage, life_stage, host_dev_stage, age_group, host_age_group | 7.9% | 11.2% |
| Treatment or growth condition | treatment, growth_condition, growth_protocol, condition, stimulus, compound, agent, treatment_protocol | 7.7% | 13.5% |
| Cell type | cell_type | 7.6% | 12.6% |
| Cell line | cell_line | 4.8% | 8.9% |
| Molecular target | chip_antibody, antibody, chip_target, antibody_target, histone_mark, target, ip_antibody, chip_antibody_target | 0.9% | 2.3% |

The values were then cleaned with a set of simple rules. Values stored as lists were reduced to their first non-empty element, and whitespace was collapsed. Values that carry no information were removed, including *NA*, *none*, *missing*, *not collected*, *not provided*, *not applicable*, *unknown*, *restricted access* and about 40 other spellings. *Sex* was mapped to a fixed set of values (*male*, *female*, *mixed*, *hermaphrodite*, *intersex*, *neuter* and *asexual*), and abbreviations such as *M* and *F*, plurals and combinations such as “*male and female*” were handled. For *age*, plain numbers with units such as *years* or “*yo*” were reduced to the number, while values such as *P56*, *E14.5*, *3 months* or *adult* were kept as written, because their meaning depends on the organism. For the molecular target, the antibody vendor, catalogue number and lot were removed, a leading “*anti-*” was dropped, and histone marks were written in one form (for example *H3K27Ac* became *H3K27ac*). All other fields were kept as written, but within each project, values that differ only in letter case were merged and shown in their most common spelling.

Finally, the run-level values were summarized for each BioProject. Because a project usually contains many samples, each field holds a list of distinct values ordered from the most to the least frequent, with at most 10 values per field, together with the number of runs in the project. Fields with no value in any run of the project are left out. Runs without a BioProject accession (about 1.3%) were skipped. Most projects end up with two to six fields. The share of runs and the share of projects with a value differ for some fields (Table 1), because a few very large surveillance projects, such as those for *SARS-CoV-2*, contain millions of runs.

### 2.3 Semantic embedding and search engine

#### 2.3.1 Embedding of BioProject records

For semantic search, each BioProject is represented by one dense vector. SRAscout uses the sentence embedding model BioBERT-mnli-snli-scinli-scitail-mednli-stsb [28], which starts from BioBERT [23] and was further trained with the Sentence-BERT approach [24] on natural language inference and sentence similarity datasets, including the clinical MedNLI. The model produces vectors of 768 dimensions. For each project, the title, name, organism and description are joined into one passage in this order, and a field is skipped when it repeats an earlier one, which often happens with the name and the title. The model reads at most 512 tokens at a time, but it was trained on short sentence pairs, so passages longer than 256 tokens are split into chunks of 256 tokens, with an overlap of 32 tokens between neighboring chunks so that sentences on a chunk border are not lost. Each chunk is embedded separately, and the vectors of all chunks are averaged into one vector for the project. In this way the whole description contributes to the embedding, while most projects, whose text is shorter than 256 tokens, are embedded as a single chunk. Projects with no text at all are not embedded. All vectors are normalized to unit length, so that cosine similarity can be computed directly. In the current release, more than 714,000 BioProjects were embedded. The same procedure is used to embed the title and abstract of an article in publication mode (Section 2.3.4).

#### 2.3.2 Storage and indexing

All data are stored in a PostgreSQL 18 database [29]. The embeddings are stored with the pgvector extension [30] and indexed with a Hierarchical Navigable Small World (HNSW) graph [31] using cosine distance. HNSW is an approximate nearest neighbor method, which finds the closest vectors among hundreds of thousands in milliseconds. For full-text search, the same four fields of each project are stored as a precomputed PostgreSQL text vector with a Generalized Inverted (GIN) Index. The fields are given different weights, with title and name weighted highest, organism second and description third, so that a word in the title counts more than the same word deep in the description.

#### 2.3.3 Search modes and hybrid search

SRAscout offers three search modes, which differ in what the user provides (Figure 1B). In text mode, the user writes free text, from a few words to a full paragraph. In publication mode, the user enters the DOI or PubMed ID of an article, and SRAscout uses its title and abstract as the query. In accession mode, the user enters one or more BioProject or SRA study accessions, and SRAscout returns these projects together with the projects most similar to them. Text mode runs a hybrid search, which combines semantic and full-text search as described below, while publication and accession modes use semantic search only (Section 2.3.4). We chose the hybrid design for text queries because each method finds what the other misses.

**Figure 1.**
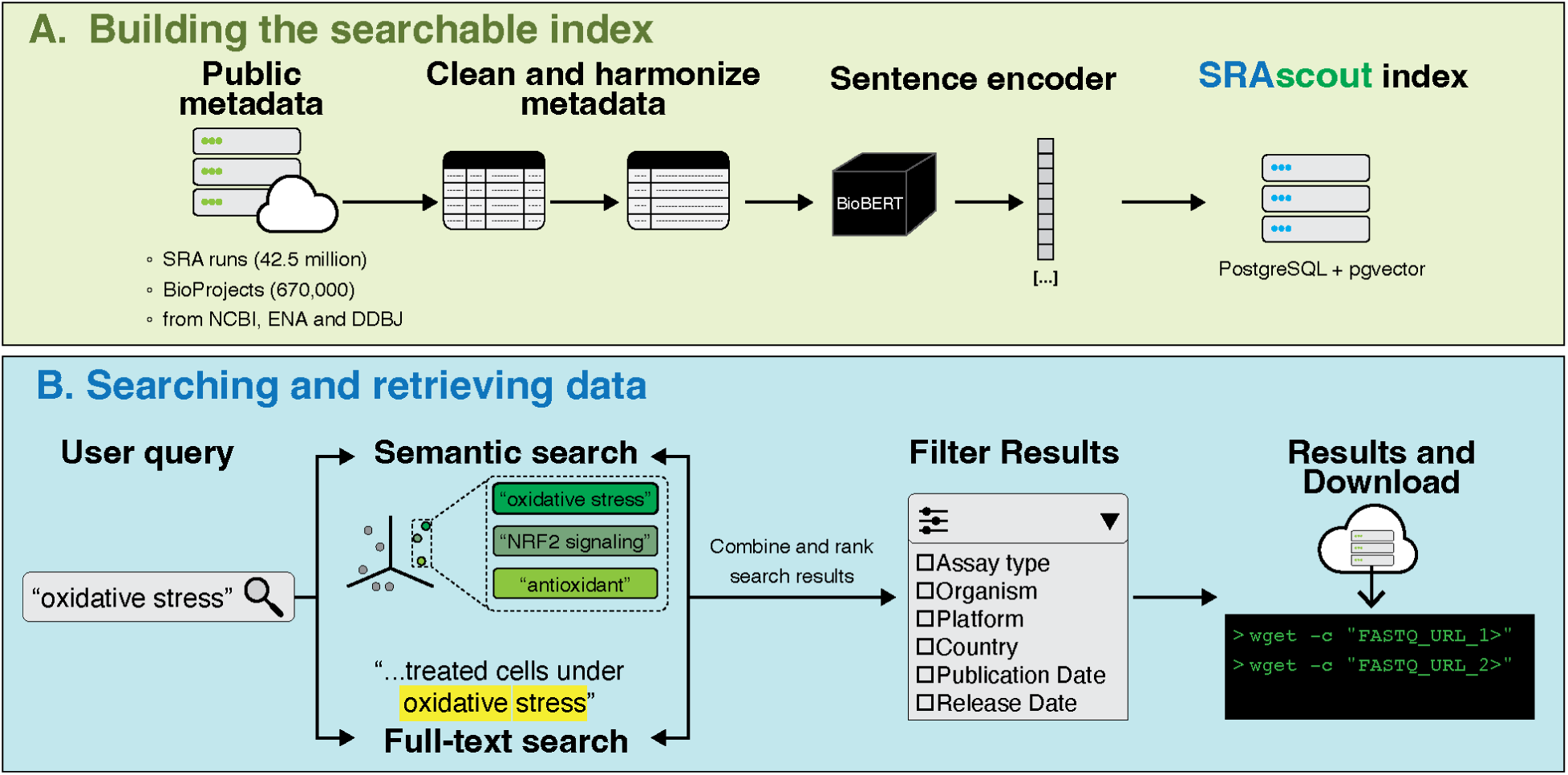
Overview of SRAscout. (**A**) Building the index. Run-level metadata of 45.2 million public runs are downloaded from the NCBI Sequence Read Archive, and project records of about 714,000 BioProjects are taken from the BioProject database shared by NCBI, ENA and DDBJ. Project titles and descriptions are cleaned, embedded with a BioBERT-based sentence embedding model and annotated with tag words extracted by GLiNER-BioMed. The free-form sample attributes of all runs are harmonized into 14 consistent fields. All data are stored in a PostgreSQL database with a vector index (HNSW) and a full-text index, and the collection is updated every week. (**B**) Searching and retrieving data. Users search in publication mode (DOI or PubMed ID), text mode (free text) or accession mode (BioProject or SRA study accessions). In text mode, a semantic search and a full-text search run together, and their ranked lists are merged by reciprocal rank fusion, while publication and accession modes use semantic search only. Results are refined with filters on organism, sample attributes, sequencing details and tag words, and users can download a harmonized metadata table and a script that fetches the sequencing files from ENA.

In the semantic part, the query is embedded with the same model and settings as the stored projects, and the 1,000 nearest projects are taken from the HNSW index. Projects with a cosine similarity below 0.5 are removed, and the rest are ranked by similarity. In the full-text part, the query is parsed with the English configuration of PostgreSQL, which reduces words to their stem (so “*mutations*” matches “*mutation*”) and ignores common words such as “*of*” and “*the*”. A project must contain every remaining word, and users can also write quoted phrases, use OR, or exclude a word with a minus sign. Matches are scored with a ranking function that is divided by the logarithm of the document length. This keeps long descriptions that repeat a word from winning, without favoring very short titles. The 1,000 best matches are kept, and among them, projects whose embedding similarity to the query is below 0.3 are removed. This step drops projects that contain the words but are mostly about something else, such as a zebrafish study that also mentions mouse samples.

The two ranked lists are merged with reciprocal rank fusion (RRF) [32]. Each project p receives the score

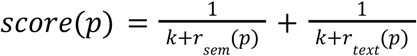

where r_sem and r_text are the ranks of the project in the semantic and full-text lists, and k = 60 which is the default value. A term is left out when the project is not in that list. Ranks are used instead of the original scores because cosine similarity and the full-text score are on different scales. With this formula, a project that ranks well in both lists comes before the top result of either list alone.

The filters selected by the user are applied to the fused list, and the results are sorted by score or by any column the user chooses. The whole search runs as a single SQL query.

#### 2.3.4 Publication and accession modes

In publication mode, the identifier entered by the user is first cleaned, so that a DOI can be given with or without its “https://doi.org/” prefix. A PubMed ID is resolved with the NCBI E-utilities [33], which return the title and abstract of the article from PubMed. A DOI is first looked up in PubMed in the same way, and when the article is not indexed there, for example a preprint or an article outside the life sciences, its metadata are retrieved from the Crossref REST API [34]. The title and abstract are joined into one passage and embedded with the same model and the same chunking as the stored projects (Section 2.3.1). When an article has no abstract, the title alone is used, and the user is informed that the search is based on the title only. If the identifier cannot be resolved, SRAscout reports this instead of running a search. The title of the resolved article is shown above the results, so users can check that the right article was found. Because a whole abstract rarely contains every word of a project description, the full-text branch would almost never match, so publication mode uses semantic search only, with the same similarity threshold of 0.5.

In accession mode, each accession is looked up in the SRAscout database. BioProject accessions from all three archives (PRJNA, PRJEB and PRJDB) and SRA study accessions (SRP, ERP and DRP) are accepted, since the BioProject records are shared by NCBI, ENA and DDBJ (Section 2.1). The matching projects are listed first. To find similar projects, SRAscout uses the stored embeddings of the given projects as the query, so no text needs to be embedded. When several accessions are given, their vectors are averaged and normalized to unit length, so that the results reflect what the projects have in common. The nearest projects are then retrieved from the HNSW index as in the semantic branch of the hybrid search, with the same threshold of 0.5, and the given projects themselves are excluded from this list. Accessions that are not found, for example projects without public sequencing runs, are reported to the user.

#### 2.3.5 Choice of thresholds

The two similarity thresholds were set by testing queries on the full set of embeddings. Relevant projects usually scored between 0.55 and 0.85. For “*soil microbiome*”, the 900th result still scored 0.70 and was on topic, and for “breast cancer tamoxifen resistance”, the results were still relevant at rank 100 (similarity 0.54). Random text scored at most about 0.55, for example “*asdf qwerty*” against a project with the very short title “*ads*”. Exact-word matches that were mostly about another topic scored as low as 0.16, while relevant ones usually scored 0.4 or higher (for “*BRCA1 mutations*”, between 0.41 and 0.56). A fixed threshold was preferred over one relative to the best score, because scores depend on the query, and broad topics score higher than specific ones.

### 2.4 Filters and metadata annotation

#### 2.4.1 Filters

The web interface offers 22 filters in three groups (Table 2). The first group describes the organism, the samples and the sequencing, the second filters by tag words, and the third describes the submitter, the size of the project and its publications. Filters can be used on their own, to browse the archive without a query, or together with a text search, in which case they are applied to the fused list of results (Section 2.3.3).

**Table 2.** Filters in the SRAscout web interface.

| Group | Filters |
|---|---|
| Organism, samples and sequencing | Organism group, organism, assay type, cell line, cell type, tissue, sex, disease, genotype or strain, treatment or condition, ChIP target, instrument, library layout, library selection, sequencing center, release date |
| Tags | Tag words, with a choice between matching all or any of the selected tags |
| Submitter, size and publications | Submitting organization, country, number of experiments, has publications, journal |

Within one filter, a project matches when it has any of the selected values, and across filters, a project must match all of them. The first group follows two different rules, because sequencing and sample information are recorded at different levels. Sequencing filters (assay type, instrument, library layout, library selection and sequencing center) must all be met by the same run, so that a search for single-end RNA-seq on a given instrument does not return a study whose RNA-seq runs and single-end runs are different experiments. Sample filters come from the harmonized attributes of Section 2.2.2 (cell line, cell type, tissue, sex, disease, genotype or strain, treatment or condition, and ChIP target) and can be met by different samples of the same project. This lets users find, for example, projects that include both male and female samples, or both treated and untreated cells.

For each list filter, the interface shows the possible values with the number of matching projects or studies, and for filters with many values it suggests matching names while the user types. Because sex was mapped to a fixed set of values, it is offered as a simple list. Results can be sorted by relevance, which is the default for text searches, or by release date, number of experiments, number of publications, accession or organism. When no query or filter is given, the most recently released projects are shown.

#### 2.4.2 Metadata annotation

Besides the fields taken from the submitted records, SRAscout adds two kinds of annotation to each BioProject. The first is the set of harmonized sample attributes described in Section 2.2.2, which provide the sample filters above. The second is a short list of tag words that names the biological entities mentioned in the title and description of the project.

The tags are extracted with GLiNER [35], a named entity recognition model that finds text spans matching a list of entity types given in plain words, and specifically with its biomedical version GLiNER-BioMed [36]. For each project, the title, the name and the full description are split into overlapping chunks in the same way as for the embeddings (Section 2.3.1), each chunk is processed separately, and the entities found in all chunks are combined. Eight types are kept as tags (organism, disease, tissue, organ, cell type, cell line, chemical or drug, and environment or habitat). Three more types (gene or protein, geographic location and sequencing technology) are recognized only so that such words are not labeled as one of the kept types, and are then discarded. GLiNER was preferred over scispaCy [37] after a comparison on 300 random projects, in which scispaCy was faster but produced many more wrong tags.

The extracted spans are then cleaned. Generic words such as cell, tissue or sample, accessions, sample identifiers, sequencer names and spans longer than five words are removed. Common words are written with only the first letter capitalized and plurals are made singular (for example mice to mouse), while acronyms, cell line names and species names keep their original form (for example HeLa and C57BL/6J). Organism names at strain level are reduced to the species, and repeated tags are removed. Finally, each tag is embedded with the same BioBERT model used for the projects, and tags whose cosine similarity to the project vector is below 0.10 are removed, with a stricter limit of 0.25 for acronyms of up to five characters. This step removes words that are only mentioned in passing, such as humans in a zebrafish study. The organism of the project is placed first, and the remaining tags follow in order of similarity, up to 20 per project. On average, a project receives about five tags, and about 3% of projects receive none.

In the tag filter, a project must have all the selected tags by default. With the switch “Match any of the tags”, users can instead find projects that have at least one of them. Because the tags are matched exactly, the same entity may appear under different names, such as Mouse and Mus musculus, and users may need to select both.

### 2.5 Web server architecture and implementation

SRAscout has three parts, a data processing pipeline, a database with a backend service, and a front-end web interface. All three are open source and are developed under the same code repository.

#### 2.5.1 Data processing pipeline

The database is built by a pipeline written in Python, which runs on a computing cluster. SRA run metadata are first collected in a local DuckDB database, filtered and then transferred to PostgreSQL. The BioProject XML file is read as a stream, so it never has to fit in memory, and the embeddings are computed on CPUs with the sentence-transformers library. Each step loads its own data into the database and builds its own indexes. Because the steps are independent, any one of them can be repeated without running the whole pipeline again.

#### 2.5.2 Database and backend

All data are stored in PostgreSQL 18 [29] database with the pgvector extension [30]. Besides the tables for projects, runs and embeddings, the database holds two materialized views that make searching fast. The first stores the full-text vector of each project (Section 2.3.2), and the second summarizes the run-level values of each SRA study, such as assay type, platform and sample country, so that run filters and their drop-down lists do not have to scan the 45.2 million runs at every search. Both views are refreshed after each data update without blocking the running service. The backend is a REST service written in Python with FastAPI [38], SQLAlchemy [39] and Psycopg, and served with gunicorn [40]. It loads the embedding model once at start-up and embeds each query in memory, so no external service is needed for semantic search. In publication mode, the backend also calls the NCBI E-utilities and, when needed, the Crossref REST API to retrieve the title and abstract of the article. Every text search, including both search branches, the rank fusion and the filters, runs as one SQL query (Section 2.3.3).

#### 2.5.3 Web interface and file downloads

The web interface is a single-page application written in TypeScript with React [41], built with Vite and styled with Ant Design and Tailwind CSS [42]. The state of every search, including the query and the selected filters, is kept in the page address, so a search can be bookmarked or shared as a link. Information about sequencing files is not stored by SRAscout. When a user opens a study, the browser requests the list of runs and files directly from the ENA Portal API, so the links are always up to date and ENA’s request limits apply to each user and not to our server.

#### 2.5.4 Updates and availability

The database is updated every week. After each update, the new projects are embedded and the materialized views, including the full-text index, are refreshed. The web server runs on a machine with 16 virtual CPUs and 32 GB of memory. SRAscout is free and open to all users without login, and it was tested on recent versions of Chrome, Firefox, Safari and Edge.

## 3. Results

### 3.1 Web server overview and usage

#### 3.1.1 Input: queries, accessions, filters

The SRAscout interface has four pages (Figure 2). The home page shows summary statistics of the indexed data and example inputs, the search page combines the search box with the filter panel and the results, the study page lists the runs and files of one SRA study, and a documentation page describes the public API. All pages are open to any user without login or registration.

**Figure 2.**
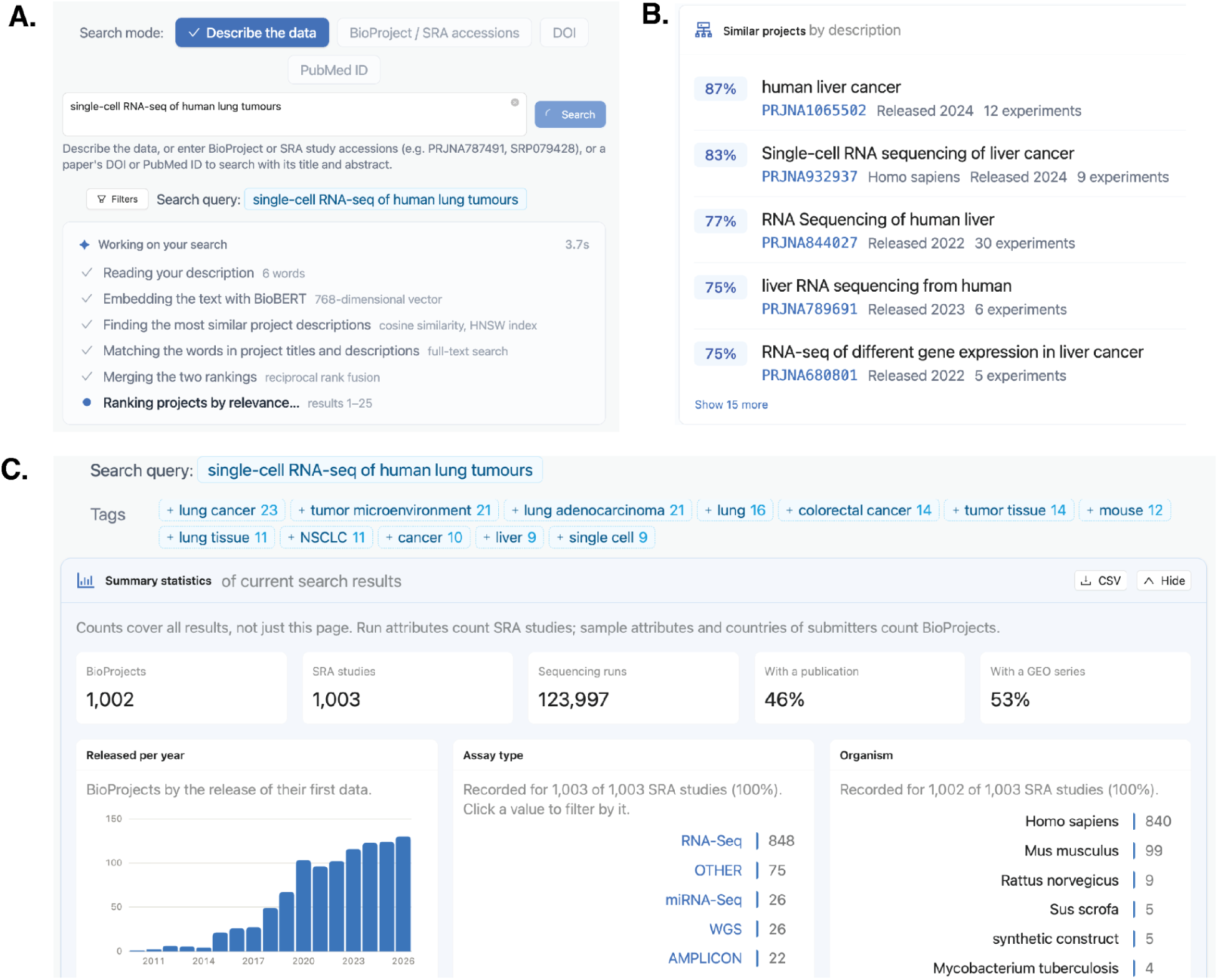
The SRAscout web interface. (**A**) The search page, with the three search modes and the filter panel, grouped into organism, samples and sequencing, tags, and submitter, size and publications. (**B**) Results of the text query “single-cell RNA-seq of human lung tumours”, with the similarity of each project to the query and a label for projects also found by full-text search. (**C**) Summary statistics of the current results, recalculated for every search, showing the number of projects, runs and publications, the release years, assay types, organisms and harmonized sample attributes, together with the tag words of the results.

Users start a search in one of three search modes (Figure 2A). In publication mode, they enter the DOI or PubMed ID of an article, and SRAscout retrieves its title and abstract and uses them as the query. This lets a researcher start from a paper they are reading, or from their own published work, and find public datasets on the same topic. In text mode, users write free text, from a few words such as “heat stress in plants” to a full paragraph such as the abstract of a planned study. In accession mode, users enter one or more BioProject or SRA study accessions from SRA, ENA or DDBJ, and SRAscout returns these projects together with the projects most similar to them. In all three modes, the input is compared with every indexed project as described in Section 2.3.

An example input for each mode can be loaded with one click, so new users can see how each mode works before entering their own.

The results can then be refined with the filters described in Section 2.4.1, which are grouped in the panel next to the results into organism, samples and sequencing, tags, and submitter, size and publications. Filters can also be used without a query, to browse the archive, for example to list all ChIP-seq projects on a given cell line. For every search, SRAscout also suggests the tag words that are most frequent among the current results, and users can add them to the search or remove them with one click to narrow or widen it. The query, the search mode and all selected filters are stored in the page address, so any search can be bookmarked, repeated later or shared with colleagues as a link.

#### 3.1.2 Output: results, project page, runs and downloads

Search results are shown as a ranked list of projects (Figure 2B). Each row gives the BioProject and SRA study accessions, the title, the organism, the number of experiments, the release date and any linked publications. For text, publication and accession searches, a Match column shows the cosine similarity between the input and each project, together with a “Words” label when the project was also found by full-text search, so users can see why each project was returned. In publication mode, the title of the resolved article is shown above the results, so users can confirm that the right article was used.

Next to the list, SRAscout shows summary statistics of the current results, such as the distribution of organisms, assay types and release years. These statistics are calculated for every search and updated whenever the query or the filters change, so users can see at a glance what kind of data their search covers and how each filter changes it. SRAscout also shows the tag words that are most frequent among the current results, which users can add to the search or remove with one click, and a network graph of these tag words, in which tags that occur together in the same projects are connected (Figure 2C). The graph helps users discover related topics, for example the tissues, diseases or treatments that are most often studied together with their query.

Selecting a project opens its project page, which brings together all the information about it. The page shows the full title and description, the harmonized sample attributes of Section 2.2.2 with the most frequent values first, the tag words, the submitting organization and the linked publications. Links lead to the original records in NCBI BioProject, ENA, GEO and PubMed. For each SRA study of the project, the study page lists its runs and, for each run, the FASTQ files with their size and MD5 checksum (Figure 2D). When ENA has no FASTQ files for a run, the files as submitted or the SRA-format file are shown instead. All links use HTTPS and can be opened directly in the browser.

Users can download the results of any search in two forms. The first is a harmonized metadata table in tab-separated format, which combines the project, sample and run information of the selected results in one file, ready to be used in an analysis or to define a cohort. The second is a shell script that downloads all the sequencing files of the selected results from ENA with wget, resumes interrupted transfers and includes the MD5 checksums of the files, so that each download can be verified. A plain list of the download links is also available for users who prefer their own download tools.

#### 3.1.3 Programmatic access (API)

SRAscout offers a read-only public API under the path /v1. It needs no registration or key and returns results in JSON. Users can search projects with the same filters as the web interface, look up a project by BioProject or SRA study accession, and list every filter with its possible values and counts. Interactive documentation, generated from the OpenAPI description of the service, lets users try each endpoint in the browser. To keep the service responsive for all users, each IP address can send one request every 5 seconds. Free-text semantic search is not yet part of the API and is planned for a future version.

### 3.2 Search performance and evaluation

#### 3.2.1 Retrieval performance

To evaluate how well SRAscout finds relevant data, we built a benchmark of 120 queries that reflect the range of questions researchers bring to public archives (Supplementary Table S1). The queries cover nine biological domains, from human disease and model organisms to plants, host-associated microbiomes and environmental samples, and five query types. Descriptive queries describe a topic in plain words, synonym queries use wording that differs from the terms submitters usually choose, identifier queries name a gene, strain, cell line or histone mark, multi-constraint queries combine several conditions, and long-text queries are paragraphs similar to a study abstract. For each query, we collected the top 20 BioProjects returned by hybrid search, by semantic search alone, by full-text search alone and by the keyword search of the NCBI BioProject database (Supplementary Table S3). All projects returned by any method were pooled, giving 5,106 pairs of a query and a project. Each pair was judged as relevant or not relevant by a large language model (Claude, Anthropic) following written criteria (Supplementary Table S2), without information on which method had retrieved the project. In total, 3,927 pairs (76.9%) were judged relevant, and every query had at least one relevant project. To validate the judgments, one author independently judged a random sample of 200 pairs, which had also been used to refine the judging criteria. The model agreed with the author on 93.0% of the pairs (Cohen’s κ = 0.82), with a precision of 91.6% and a recall of 99.3% against the author’s labels. Most disagreements were projects that the model judged relevant and the author did not (13 of 14), so the model was slightly more lenient than the author. We measured recall at 20, the share of the relevant projects for a query that appear among its top 20 results, and averaged it over all queries.

Hybrid search retrieved 60.3% of the relevant projects among the top 20 results, compared with 56.1% for semantic search alone, 30.6% for full-text search alone and 23.6% for the NCBI BioProject search (Figure 3, Supplementary Table S4). The semantic component made the largest difference for queries whose wording differs from the project descriptions. For synonym queries, hybrid search reached 66.8% compared with 18.1% for NCBI. For example, the query “nerve cells dying in neurodegeneration” found all relevant projects with hybrid search and none with keyword search. Long-text queries were answered almost entirely by the semantic component (96.7% for semantic search compared with 4.7% for full-text search), since a paragraph rarely matches a project description word for word. Multi-constraint queries also benefited strongly, with 83.0% for hybrid search compared with 13.4% for NCBI. For identifier queries, full-text search alone (38.8%) performed better than semantic search alone (31.8%), and hybrid search kept this strength and slightly improved on it (40.5%), showing that merging the two lists preserves exact matches on gene names, strains and cell lines. Hybrid search was the best or close to the best method in all nine domains (Supplementary Table S4).

**Figure 3.**
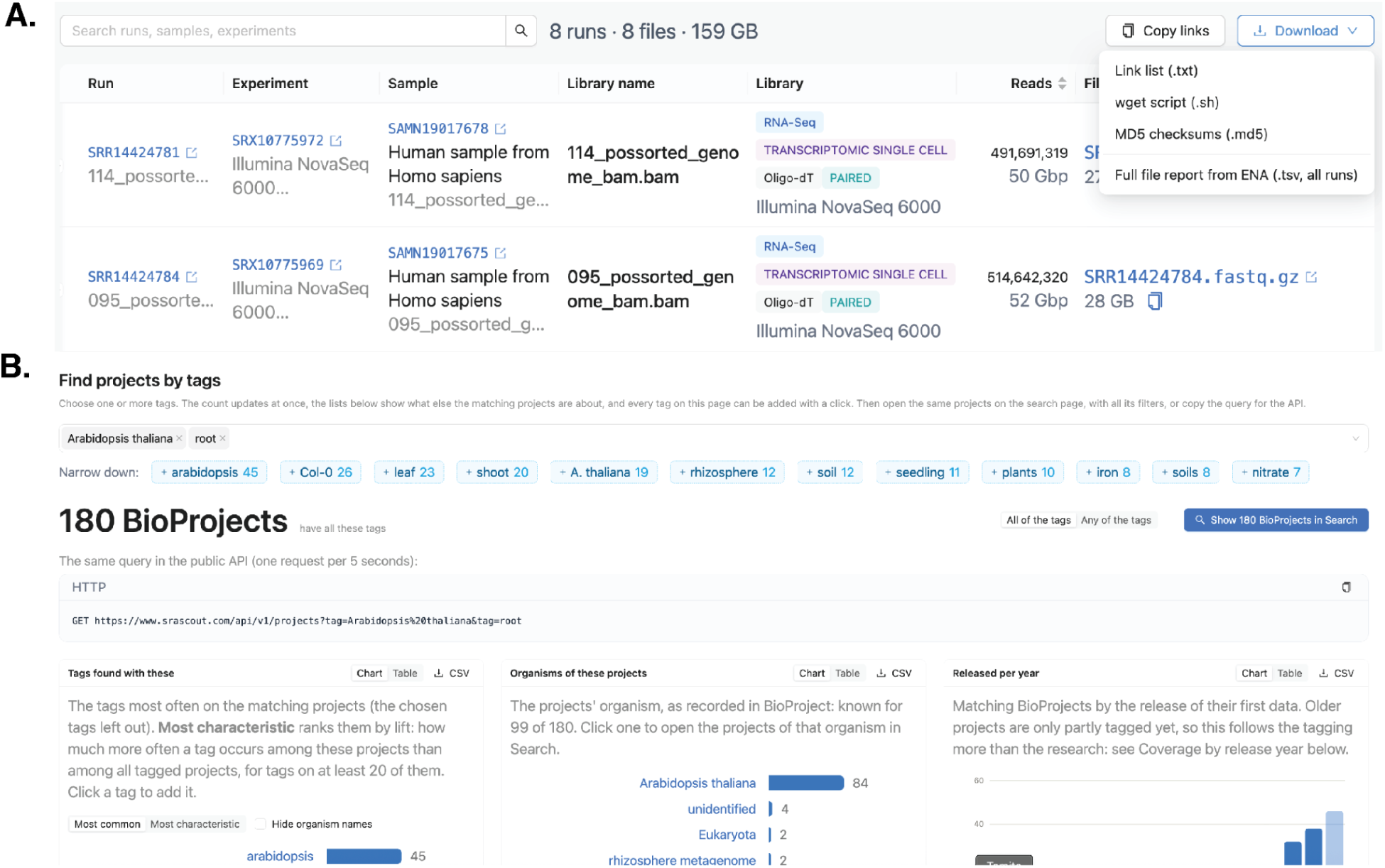
Project details, data download and tag-based search in SRAscout. (A) The study page of *PRJNA727404*, showing the title and description of the project, its harmonized sample attributes, tag words, submitting organization and linked publications, with links to NCBI BioProject, ENA, GEO and PubMed. The runs of the study are listed with their FASTQ files, sizes and MD5 checksums retrieved from the ENA Portal API, together with the options to download the harmonized metadata table, the download script and the list of download links. (B) Finding BioProjects with tag words. The tags *arabidopsis thaliana* and *root* are selected in the tag filter, requiring all selected tags, and the matching projects are listed. The tags most frequent among the results can be added to or removed from the search with one click. The network graph connects tags that occur together in the same projects.

SRAscout also reaches projects that are invisible to publication-based search. Of the 1,850 distinct relevant projects found by hybrid search, 63.6% have no linked publication and 59.2% have no GEO series, so most of them could not have been found with tools built on PubMed or GEO. On the production server, a hybrid search took a median of 0.90 seconds over the 120 benchmark queries (range 0.28 to 4.39 seconds), of which a median of 98 ms was spent on embedding the query.

#### 3.2.2 Accuracy of harmonized attributes

To check that the harmonized sample attributes can be trusted as filters, we drew a random sample of 140 harmonized values, 10 for each of the 14 fields in Table 1 (Supplementary Table S5). For each value, one author compared the harmonized values with the original attributes written by the submitters and labeled them as correct, partly correct or incorrect. A value was counted as partly correct when it was right but incomplete, and as incorrect when it did not match the original meaning. Of the 140 values, 139 were correct (99.3%, 95% confidence interval 96.1% to 99.9%), one was partly correct and none was incorrect. The partly correct value was in the host field. These results show that the cleaning rules keep the meaning of the submitted values. However, harmonization cannot correct errors made by the submitters themselves. For example, some projects record a cell line or a diagnosis in the cell type field, and such values remain in the filters as they were written. With 10 values per field, the sample also cannot rule out rare errors in individual fields.

### 3.3 Feature comparison with existing tools

Table 3 compares SRAscout with existing tools for finding public sequencing datasets. Archive interfaces such as NCBI SRA, the ENA Browser and DDBJ Search cover all data but rely on keyword search over submitted fields. MetaSRA and SRAgent harmonize sample metadata with ontologies, but MetaSRA covers only human and mouse RNA-seq and ChIP-seq samples up to 2020, and SRAgent is a command-line tool whose dataset search is limited to single-cell RNA-seq. GEOfinder and POE support semantic search but are limited to GEO, and POE requires a linked publication. SRAscout is the only tool in this comparison that combines free-text semantic search over all SRA BioProjects with filters on harmonized sample attributes and direct access to the sequencing files.

**Table 3.**
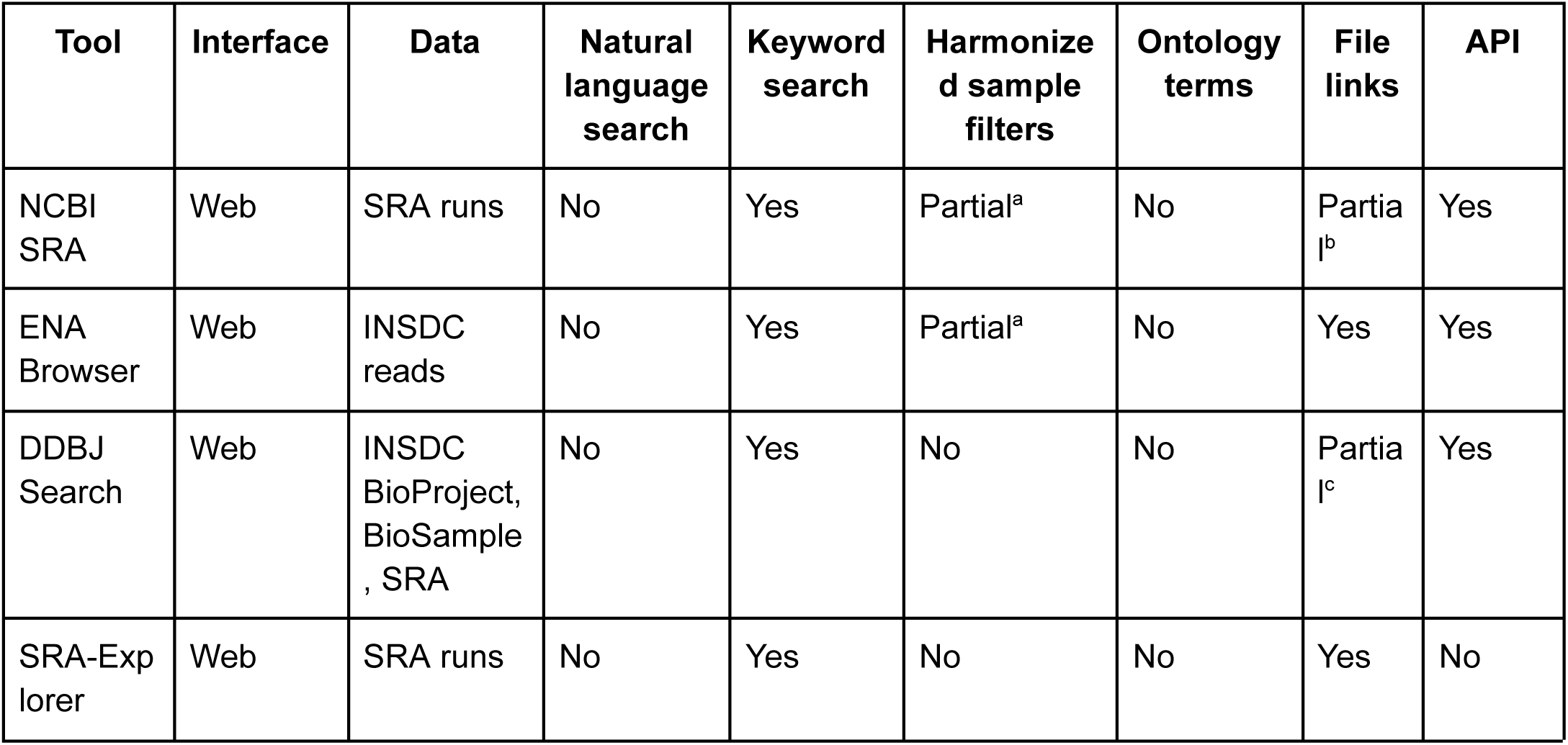
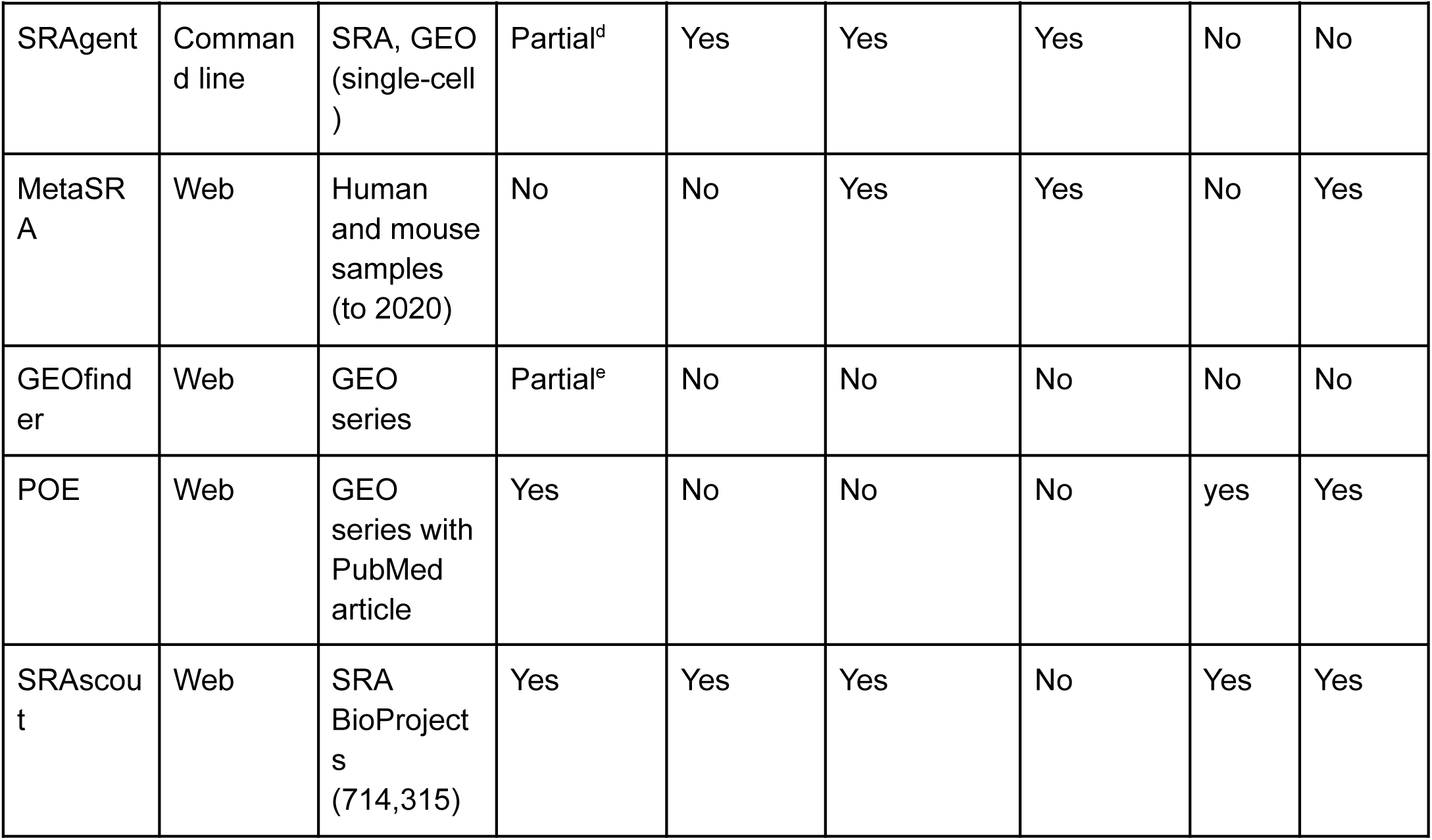
Features of SRAscout and of existing tools for finding public sequencing datasets.

| **Tool** | **Interface** | **Data** | **Natural language search** | **Keyword search** | **Harmonized sample filters** | **Ontology terms** | **File links** | **API** |
|---|---|---|---|---|---|---|---|---|
| NCBI SRA | Web | SRA runs | No | Yes | Partial<sup>a</sup> | No | Partial<sup>b</sup> | Yes |
| ENA Browser | Web | INSDC reads | No | Yes | Partial<sup>a</sup> | No | Yes | Yes |
| DDBJ Search | Web | INSDC BioProject, BioSample, SRA | No | Yes | No | No | Partial<sup>c</sup> | Yes |
| SRA-Explorer | Web | SRA runs | No | Yes | No | No | Yes | No |
| SRAgent | Command line | SRA, GEO (single-cell) | Partial<sup>d</sup> | Yes | Yes | Yes | No | No |
| MetaSRA | Web | Human and mouse samples (to 2020) | No | No | Yes | Yes | No | Yes |
| GEOfinder | Web | GEO series | Partial<sup>e</sup> | No | No | No | No | No |
| POE | Web | GEO series with PubMed article | Yes | No | No | No | yes | Yes |
| SRAscout | Web | SRA BioProjects (714,315) | Yes | Yes | Yes | No | Yes | Yes |

Yes, available; Partial, available with limitations; No, not available. INSDC, International Nucleotide Sequence Database Collaboration; API, application programming interface. ^a^Facets on submitted fields such as organism, library strategy and platform. ^b^Run lists from the SRA Run Selector; files are obtained with the SRA Toolkit. ^c^Files on the DDBJ mirror, which does not hold all SRA runs. ^d^An LLM agent writes Entrez keyword queries; results are not ranked by similarity. ^e^Search by example GEO series accessions, not by free text.

## 4. Discussion

Public sequencing archives hold an enormous amount of data that could be reused, but finding the right studies is still one of the main challenges. SRAscout addresses this problem by combining three things in one public web server. It indexes the BioProjects behind all public SRA runs, it lets users search them in plain language through a hybrid of semantic and full-text search, and it lets them narrow the results with filters on harmonized sample attributes, sequencing details and tag words. Each result then leads directly to the runs and their sequencing files in ENA, so the path from a research question to downloaded data takes a few steps in the browser.

Compared with our previous work, POE, SRAscout covers a much larger and more varied part of the public data. POE reaches only GEO datasets with a PubMed publication, because it builds its search on the text of the linked articles. SRAscout instead uses the text of the BioProject records themselves, so it also reaches projects that were deposited directly in SRA or ENA, projects from all organisms and assay types, and recent projects that have not yet been published. As a result, SRAscout indexes more than 42 million sequencing runs, about four times the roughly 10 million samples covered by POE. Compared with other tools (Table 3), SRAscout is, to our knowledge, the only public web server that combines natural language search over all SRA BioProjects with filters on harmonized sample attributes (tissues, cell types, cell lines) and direct access to the sequencing files.

The hybrid design was a deliberate choice. Semantic search alone finds studies that describe the same biology in other words, but it handles exact identifiers poorly and tends to rank very short texts too high. Full-text search alone finds exact terms such as gene names and strain identifiers, but it misses studies that use different words. Merging the two ranked lists with reciprocal rank fusion keeps the strengths of both, and the results show users whether each project was found by meaning, by exact words, or by both.

The harmonization of sample attributes follows simple and transparent rules instead of a trained model. This makes the process fast, easy to repeat at every update and easy to inspect, since every value in a filter can be traced back to the key and the text that the submitter wrote. The filters also respect how the data are organized. Sequencing filters must be met by the same run, while sample filters can be met by different samples, which matches the questions researchers usually ask, such as finding projects that include both treated and untreated samples sequenced with the same method.

SRAscout also has limitations. First, projects with very short titles and no description carry little information, and they can still receive high similarity scores for many unrelated queries. The fusion with full-text search reduces this effect, but it does not remove it for queries without exact word matches. Second, the full-text branch requires every word of the query, so long descriptive queries rely mostly on the semantic branch. Third, the harmonized attributes are cleaned but not mapped to ontologies, so synonyms such as different names for the same tissue remain separate values, and several fields, such as disease state, are filled in only for a small share of projects because submitters rarely provide them. Fourth, SRAscout indexes only public metadata. Human sequencing data from clinical settings, such as diagnostic gene panels on tissue and liquid biopsies [43] or hereditary cancer risk testing [44], are usually held in controlled-access and are outside its scope.

In summary, SRAscout makes the metadata of public sequencing data easier to search, filter and reuse. By connecting natural language search with structured filters and direct file access, it helps researchers find relevant datasets for new analyses, build cohorts for meta-analyses and reuse existing data instead of generating it again.

## Data Availability

SRAscout does not host or store raw omics data. Instead, it indexes structured metadata and provides direct external links to corresponding entries in SRA, GEO, PubMed, and ENA.

SRAscout is a freely available web application, accessible at www.srascout.com.

## Code availability

Code for the data processing pipeline, the backend and the web interface is available at https://github.com/Georgakopoulos-Soares-lab/sra_explorer.

## Supporting information

Supplementary_benchmark

## References

1. Leinonen R, Sugawara H, Shumway M, International Nucleotide Sequence Database Collaboration. The sequence read archive. Nucleic Acids Res. 2011;39: D19–21.

2. Leinonen R, Akhtar R, Birney E, Bower L, Cerdeno-Tárraga A, Cheng Y, et al. The European Nucleotide Archive. Nucleic Acids Res. 2011;39: D28–31.

3. Grigoriadis D, Perdikopanis N, Georgakilas GK, Hatzigeorgiou AG. DeepTSS: multi-branch convolutional neural network for transcription start site identification from CAGE data. BMC Bioinformatics. 2022;23: 395.

4. Perdikopanis N, Georgakilas GK, Grigoriadis D, Pierros V, Kavakiotis I, Alexiou P, et al. DIANA-miRGen v4: indexing promoters and regulators for more than 1500 microRNAs. Nucleic Acids Res. 2021;49: D151–D159.

5. Tsifintaris M, Grigoriadis D, Sitmalidis M, Repanas P, Lazarashvili EK, Anastasiadi C, et al. ASTRA: a comprehensive resource of stress-induced transcriptional activity in human cell lines. Nucleic Acids Res. 2026;54: D938–D948.

6. Wilkinson MD, Dumontier M, Aalbersberg IJJ, Appleton G, Axton M, Baak A, et al. The FAIR Guiding Principles for scientific data management and stewardship. Sci Data. 2016;3: 160018.

7. Gonçalves RS, Musen MA. The variable quality of metadata about biological samples used in biomedical experiments. Sci Data. 2019;6: 190021.

8. Bernstein MN, Doan A, Dewey CN. MetaSRA: normalized human sample-specific metadata for the Sequence Read Archive. Bioinformatics. 2017;33: 2914–2923.

9. Zhu Y, Stephens RM, Meltzer PS, Davis SR. SRAdb: query and use public next-generation sequencing data from within R. BMC Bioinformatics. 2013;14: 19.

10. Choudhary S. pysradb: A Python package to query next-generation sequencing metadata and data from NCBI Sequence Read Archive. F1000Res. 2019;8: 532.

11. Chao H, Li Z, Chen D, Chen M. iSeq: an integrated tool to fetch public sequencing data. Bioinformatics. 2024;40: btae641.

12. Patel H, Garcia MU, Talbot A, Sateesh_Peri, Beber ME, Joshi E, et al. nf-core/fetchngs: nf-core/fetchngs v1.12.0 - Titanium Platypus. Zenodo; 2024. doi:10.5281/ZENODO.10728509

13. Ewels P. SRA Explorer. [cited 14 Nov 2026]. Available: https://sra-explorer.info/#

14. Fukuda A, Kodama Y, Mashima J, Fujisawa T, Ogasawara O. DDBJ update: streamlining submission and access of human data. Nucleic Acids Res. 2021;49: D71–D75.

15. Hicks P, Valtadoros LE, Mancuso CA, Alquadoomi F, Johnson KA, Sundar S, et al. MetaHQ: Harmonized, high-quality metadata annotations of public omics samples and studies. 2026. Available: http://arxiv.org/abs/2602.07805

16. Youngblut ND, Carpenter C, Prashar J. scBaseCount: an AI agent-curated, uniformly processed, and continually expanding single cell data repository. bioRxiv. doi:10.1101/2025.02.27.640494

17. Thalhath N. srake. Github; Available: https://github.com/nishad/srake

18. Wilks C, Zheng SC, Chen FY, Charles R, Solomon B, Ling JP, et al. recount3: summaries and queries for large-scale RNA-seq expression and splicing. Genome Biol. 2021;22: 323.

19. Lachmann A, Torre D, Keenan AB, Jagodnik KM, Lee HJ, Wang L, et al. Massive mining of publicly available RNA-seq data from human and mouse. Nat Commun. 2018;9: 1366.

20. Perez-Riverol Y, Bai M, da Veiga Leprevost F, Squizzato S, Park YM, Haug K, et al. Discovering and linking public omics data sets using the Omics Discovery Index. Nat Biotechnol. 2017;35: 406–409.

21. Chikhi R, Lemane T, Loll-Krippleber R, Montoliu-Nerin M, Raffestin B, Camargo AP, et al. Logan: Planetary-Scale Genome Assembly Surveys Life’s Diversity. bioRxiv. 2025. p. 2024.07.30.605881. doi:10.1101/2024.07.30.605881

22. Karasikov M, Mustafa H, Danciu D, Kulkov O, Zimmermann M, Barber C, et al. Efficient and accurate search in petabase-scale sequence repositories. Nature. 2025;647: 1036–1044.

23. Lee J, Yoon W, Kim S, Kim D, Kim S, So CH, et al. BioBERT: a pre-trained biomedical language representation model for biomedical text mining. Bioinformatics. 2020;36: 1234–1240.

24. Reimers N, Gurevych I. Sentence-BERT: Sentence embeddings using Siamese BERT-networks. arXiv [cs.CL]. 2019. doi:10.48550/arXiv.1908.10084

25. Brown GS, Wengler J, Fabelico AJS, Muir A, Tubbs A, Warren A, et al. Using semantic search to find publicly available gene-expression datasets. Bioinformatics. 2026;42. doi:10.1093/bioinformatics/btag053

26. Grigoriadis D, Tsifintaris M, Giannakakis A, Pavlopoulos GA, Perdikopanis N. Public Omics Explorer (POE): Enabling integrative semantic search across GEO omics datasets based on PubMed publications. Comput Struct Biotechnol J. 2025;27: 4802–4812.

27. Barrett T, Clark K, Gevorgyan R, Gorelenkov V, Gribov E, Karsch-Mizrachi I, et al. BioProject and BioSample databases at NCBI: facilitating capture and organization of metadata. Nucleic Acids Res. 2012;40: D57–63.

28. pritamdeka/BioBERT-mnli-snli-scinli-scitail-mednli-stsb · Hugging Face. [cited 14 Nov 2026]. Available: https://huggingface.co/pritamdeka/BioBERT-mnli-snli-scinli-scitail-mednli-stsb

29. PostgreSQL. In: PostgreSQL [Internet]. 14 Nov 2026 [cited 14 Nov 2026]. Available: https://www.postgresql.org/

30. pgvector: Open-source vector similarity search for Postgres. Github; Available: https://github.com/pgvector/pgvector

31. Malkov YA, Yashunin DA. Efficient and robust approximate nearest neighbor search using hierarchical Navigable Small World graphs. IEEE Trans Pattern Anal Mach Intell. 2020;42: 824–836.

32. Cormack GV, Clarke CLA, Buettcher S. Reciprocal rank fusion outperforms condorcet and individual rank learning methods. Proceedings of the 32nd international ACM SIGIR conference on Research and development in information retrieval. New York, NY, USA: ACM; 2009. doi:10.1145/1571941.1572114

33. Sayers E. A General Introduction to the E-utilities. Entrez Programming Utilities Help Bethesda (MD): National Center for Biotechnology Information (US). 2010.

34. Lammey R. Using the Crossref Metadata API to explore publisher content. Sci Ed. 2016;3: 109–111.

35. Zaratiana U, Tomeh N, Holat P, Charnois T. GLiNER: Generalist Model for Named Entity Recognition using Bidirectional Transformer. Proceedings of the 2024 Conference of the North American Chapter of the Association for Computational Linguistics: Human Language Technologies (Volume 1: Long Papers). 2024. pp. 5364–5376.

36. Yazdani A, Stepanov I, Teodoro D. GLiNER-BioMed: A Suite of Efficient Models for Open Biomedical Named Entity Recognition. 2025. Available: http://arxiv.org/abs/2504.00676

37. Neumann M, King D, Beltagy I, Ammar W. ScispaCy: Fast and robust models for biomedical natural language processing. Proceedings of the 18th BioNLP Workshop and Shared Task. Stroudsburg, PA, USA: Association for Computational Linguistics; 2019. doi:10.18653/v1/w19-5034

38. FastAPI - FastAPI. [cited 14 Nov 2026]. Available: https://fastapi.tiangolo.com/

39. SQLAlchemy. [cited 14 Nov 2026]. Available: https://www.sqlalchemy.org/

40. Gunicorn - Python WSGI HTTP Server for UNIX. [cited 14 Nov 2026]. Available: https://gunicorn.org/

41. React. [cited 14 Nov 2026]. Available: https://react.dev/

42. Rapidly build modern websites without ever leaving your HTML. [cited 14 Nov 2026]. Available: https://tailwindcss.com/

43. Meintani A, Ozdogan M, Touroutoglou N, Papazisis K, Boukovinas I, Bilir C, et al. Comprehensive evaluation of a 1021-gene panel in FFPE and liquid biopsy for analytical and clinical use. Int J Mol Sci. 2025;26: 5930.

44. Tsoulos N, Papadopoulou E, Agiannitopoulos K, Grigoriadis D, Tsaousis GN, Bouzarelou D, et al. Polygenic Risk Score (PRS) combined with NGS panel testing increases accuracy in hereditary breast cancer risk estimation. Diagnostics (Basel). 2024;14: 1826.

